# PROTACable is an Integrative Computational Pipeline of 3-D Modeling and Deep Learning to Automate the De Novo Design of PROTACs

**DOI:** 10.1101/2023.11.20.567951

**Authors:** Hazem Mslati, Francesco Gentile, Mohit Pandey, Fuqiang Ban, Artem Cherkasov

## Abstract

Proteolysis-targeting chimeras (PROTACs) that engages two biological targets at once is a promising technology in degrading clinically relevant protein targets. Since factors that influence the biological activities of PROTACs are more complex than those of a small molecule drug, we explored a combination of computational chemistry and deep learning strategies to forecast PROTAC activity and enable automated design. A new method named PROTACable was developed for de novo design of PROTACs, which includes a robust 3-D modeling workflow to model PROTAC ternary complexes using a library of E3 ligase and linker and an SE(3)-equivariant graph transformer network to predict the activity of newly designed PROTACs. PROTACable is available at https://github.com/giaguaro/PROTACable/.

## Introduction

Targeted therapies in cancer treatment often rely on small-molecule inhibitors (SMIs) to block harmful proteins. While effective, this conventional approach face a number of challenges that extend beyond the potential for off-target toxicity, including the requirement of high selectivity, the potential for reduced effectiveness due to point mutations, and the risk that the inhibitor-protein complex could trigger oncogenic pathways^1–3^. Protein degradation, on the other hand, is a natural process of proteins’ turnover. Synthetic proteolysis targeting chimeras (PROTACs) are a novel class of heterobifunctional drugs that harness this natural process in order to selectively degrade biological targets of interest intracellularly using host cell’s E3 ubiquitin ligases. PROTACs have recently attracted particular interest in drugging “undruggable targets” characterized with binding sites that are shallow, possess promiscuous conformations, or are susceptible to mutations – thus intractable with small-molecule inhibitors^4^.

Crafting a PROTAC requires three chemical moieties: 1) a warhead that selectively binds the to-be-degraded protein of interest (POI)^5^; 2) a ligand that binds to the host cell’s E3 ubiquitin ligase machinery; and 3) a linker that connects the two POI and E3 ligands^6^. The resulting molecule then triggers the formation of a ternary complex by baiting POI within a close proximity to a ubiquitin ligase (the cell’s *garbage* disposal system), where it gets marked for post-translational degradation^3,6^. The predominant approach to PROTAC design utilizes known ligands or their modified versions for common E3 ligases such as Von-Hippel-Lindau (VHL), Cereblon (CRBN), Bruton’s tyrosine kinase (BTK), murine double minute 2 (MDM2), Ubiquitin Protein Ligase E3 Component N-Recognin 1 (UBR1), and cell inhibitor of apoptosis protein (cIAP) ubiquitin ligases, among others^7^.

However, unlike SMIs, designing PROTACs presents significant challenges, including the complex structure-activity relationship optimization, the synthetic difficulty of heterobifunctional molecules, and the need for a detailed understanding of ternary complex formation to enhance the drug-like properties and pharmacokinetics of these large molecular weight compounds^8,9^. For instance, a link between E3 ligase choice and PROTAC activity has been observed^8,10^. E3 ligase bioavailability is influenced by tissue expression, while varying thermodynamic cooperativity is observed with POI-E3 binding and PROTAC molecule saturation^8^. Similarly, the design of the linker is also crucial to ensure stability and specificity of the ternary complex^10,11^.

Recently, a range of computational methods have emerged to help the discovery of innovative PROTACs^5,11–18^. However, these methods often require substantial resources and an intricate grasp of the target’s tertiary or quaternary structure. Moreover, earlier research^19–26^ predominantly validated these methods only on active crystallized PROTACs, thereby curbing their broader applicability. On the other hand, commercial tools^5,27^, which offers enumerating solutions for ternary conformations, require prepared and known components of the ternary complex. In fact, pinpointing the native ternary complex remains a challenge due to the constraints of physics-based scoring functions^28^, particularly concerning protein-protein interactions. Despite this, available *in silico* tools are often specialized in tackling one aspect at a time in the design of a functional PROTAC such as the selection of a suitable E3 ligase^29^, the optimization of the linker^30^, or prediction of affinity^18^. Consequently, there exists a need for establishing a pipeline that integrates of the multifaceted criteria for PROTAC formulation *in* silico.

This work implemented a widely applicable computational pipeline for the *de novo* design of PROTACs. We first developed and benchmarked a modeling protocol against the available crystallized PROTAC ternary complexes by comparing the structures’ alpha-carbon (Cα-RMSD) to the native ternary proteins, and the ligand (L-RMSD) to the native PROTACs; then employed the workflow to model the 1,236 entries of the PROTACpedia database^31^ into 3-D ternary complexes. Given these structurally sound complexes, an SE(3)-equivariant graph transformer network was trained using 3-D atomic representation to score filtered ternary complex poses based on the likelihood of *in vitro* activity. The 3-D equivariance which is characteristic of the SE(3)-transformer enables discerning critical structural attributes, irrespective of the input structure’s spatial orientation or positioning^32^. In addition, we packaged the modelled E3 ligase and linker library in a workflow that streamlines the design of novel PROTACs.

Our strategy has demonstrated state-of-the-art performances of both ternary structure prediction and activity classification for efficient and automated design of novel PROTACs.

### Related works

This work focuses on the modeling of the ternary complex and its validation through crystallographic benchmarking, the comprehensive design of PROTAC molecules, and the prediction of PROTAC activity.

Recently Liao et al.^12^, Zaidman et al.^14^, and Weng et al.^13^ leverage computational tools to model ternary complexes. Liao et al. proposes a molecular dynamics (MD)-based strategy and heating-accelerated pose change (HOPAC) rescoring method, which requires high computational resources and manual intervention^12^. Zaidman et al. provide an automated pipeline (PRosettaC), particularly at the linker insertion stage, although their method requires expert tuning based on selected atom identifiers within the PROTAC SMILES string^14^. Weng et al.’s approach necessitates of an established holo-form of protein complexes and user intervention for pose selection^13^. Recent works also include PROTAC-RL pipeline by Zheng et al.^17^ The reinforcement learning (RL) agent in PROTAC-RL uses a ‘prior’ network (Proformer), trained on small molecules to guide selection of desirable pharmacokinetic properties (PK) of linkers given input POI and E3 ligase ligands. The PROTAC-RL follows PRosettaC in modeling the ternary complex. PROTAC-INVENT by Li et al.^33^, on the other hand, transforms 2-D RL-generated linkers into 3-D conformers and reconstructs PROTAC confirmation given a starting POI-ligand (POI^Lig^) and E3-ligand (E3^Lig^) pairs for guiding a constrained MD.

In the context of PROTAC design and activity prediction, Nori et al.^16^ proposes PROTAC-Design, a generative model coupled with RL which sample new molecules from a prior representative of trained datapoints from PROTAC-db^34^. Li et al.^18^ developed DeepPROTACs, a deep learning model for predicting the efficiency of PROTACs using graph convolutional networks (GCNs) with a long-short term memory (LSTM) layer to incorporate structural and sequence information. The model separately considers the structures of the protein pocket, the E3 ligase pocket, and the PROTACs, achieving an accuracy of up to 78%^18^.

### Our contributions

- Our modelling approach achieves superior performance in recapturing crystallized ternary complexes compared with available open-source and commercial workflows.
- We introduce the first end-to-end PROTAC design tool while leveraging E3 ligase library and experimentally verified linkers for potential candidates. This is complemented by a high-performing SE(3)-equivariant graph transformer that predicts ternary activity.

In contrast to several other works, our automated pipeline only needs a target POI and its corresponding cognate ligand, operates in full 3-D atomic resolution, and considers inactive PROTACs in activity predictions.

## Results and Discussion

### Modeling the ternary complexes

A computational workflow was developed and validated to model ternary complexes starting from the individual structures of the PROTAC, the POI, and the E3 ligase of known PROTACs from the PROTACpedia database^31^ (**Figure 1**). Each individual PROTACpedia entry was thus deconstructed and anchoring sites were annotated with a “radioactive” tag on their terminal atoms. The modeling procedure was then applied to all the 1,236 entries.

**Figure 1.**
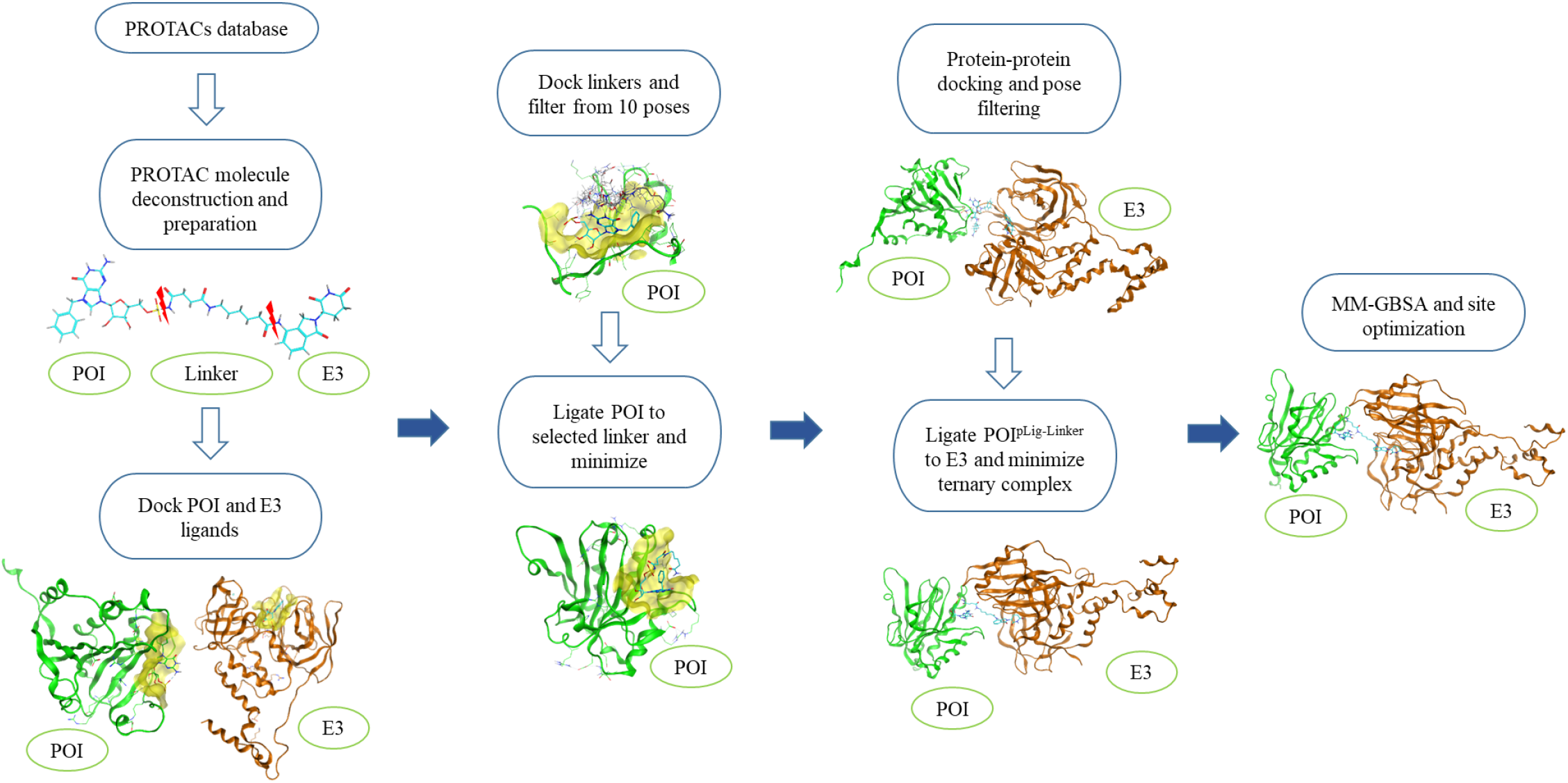
The workflow developed to model PROTAC ternary complexes. The ligands are shown in cyan, the POI in dark green, and the E3 in dark orange. The pipeline consists of the following steps: 1) A given 2-D PROTAC is deconstructed into its building blocks: the POI cognate ligand, the linker, and the E3 cognate ligand. 2) Each cognate ligand – POI^Lig^ and E3^Lig^ – is docked separately into the corresponding pre-identified pockets on POI and E3, respectively. 3) Up to 10 linker conformations are sampled from docking the linker in the POI pocket in the presence of the POI^pLig^ docked in step 2. The linker corresponding to the shortest distance between the linker anchor point and the POI^Lig^ anchoring point is selected and retained. 4) The linker is ligated from its attachment site to the POI^pLig^ anchoring site. A system energy re-equilibration is performed through minimization of the complex afterwards. 5) Unconstrained protein-protein docking between POI^pLig-Linker^ complex and the corresponding E3^eLig^ complex takes place resulting in 100 poses. The poses are filtered down to 20 PROTAC-productive conformations using an in-house developed algorithm (see **Methods**). 6) The final ligation between the POI^pLig-Linker^ and the E3^eLig^ is accomplished using an automated PDB file text-manipulation script. This is ensued by another round of minimization for system re-equilibration. 7) Molecular Mechanics - Generalized Born Solvent Area (MM-GBSA) rescoring, binding-site optimization and energy calculations are performed for pocket minimization and rescoring for ranking the 20 ternary complexes.

The pipeline was validated on 15 complexes obtained from the RCSB Protein Data Bank^35^, representing crystallized PROTACs to date of the publication’s writing with PDB IDs: 5T35, 6BN7, 6BOY, 6HAX, 6HAY, 6HR2, 6SIS, 6W7O, 6W8I, 6ZHC, 7JTO, 7JTP, 7KHH, 7PI4, and 7Q2J^19–24,26,36^ (**Figure S1**). The top 20 modeling solutions from each complex were evaluated for protein-to-protein alpha carbon RMSD (Cα-RMSD) and ligand-to-ligand (L-RMSD) against the crystallized complex. The lowest RMSD solutions for the eight popularly benchmarked ternary complexes^5,12–14^ are shown in **Figure 2**.

**Figure 2.**
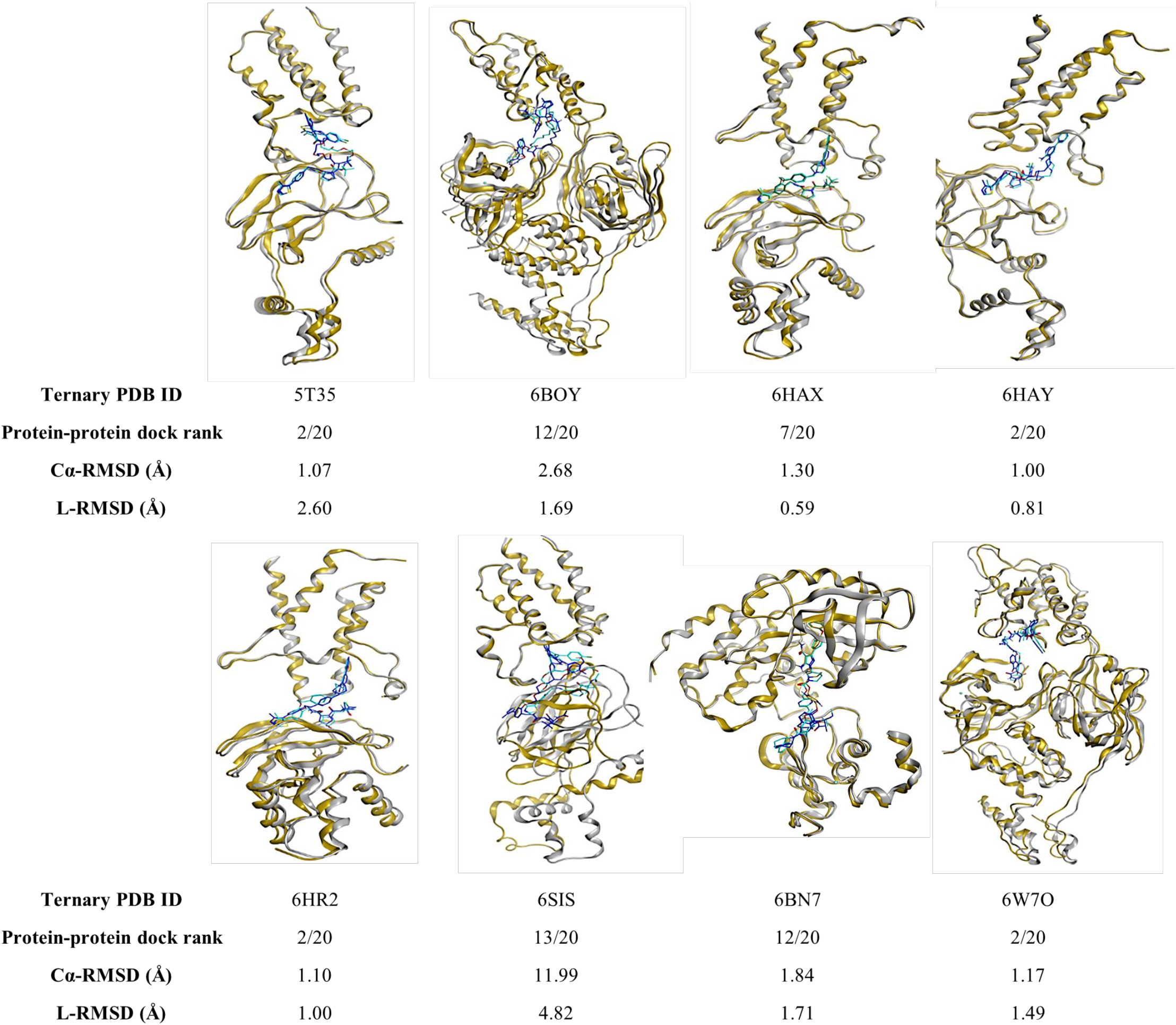
Benchmarking of the pipeline on eight crystallized ternary complexes. Native ternary proteins are colored in grey while their respective native PROTAC poses are colored in blue. Modelled ternary proteins are colored in gold while respective modelled PROTAC poses are colored in cyan. Below each complex, the solution rank (out of 20) is shown as well as the Cα-RMSD and the PROTAC L-RMSD between the modeled and the experimental structure.

Considering 10 Ǻ between predicted and crystal protein-protein complexes as RMSD threshold for docking success^15,37^, our method accurately recapitulated 9 out 15 structures, while 3 were borderline unacceptable (slightly above 10 Ǻ), and 3 were rejected. Considering the PROTAC ligands, the RMSD was below 2.0 Ǻ in 8 cases^38^. Notably, several PROTAC ligands demonstrated good ligand L-RMSD despite suboptimal Cα-RMSD – such as with the case of 6W8I, 7JTO and 7Q2J (**Figure 2** and **Table 1**).

**Table 1.** Comparison of our method with existing methodologies in literature for modeling PROTAC ternary complexes.

| Study | Ranking method |  | 5T35 | 6BN7 | 6BOY | 6HAX | 6HAY | 6HR2 | 6SIS | 6W7O |
| --- | --- | --- | --- | --- | --- | --- | --- | --- | --- | --- |
| This method | MM-GBSA<br>(crystal set) | **Rank<sup>a</sup>** | 2/20 | 5/20 | 4/20 | 1/20 | 3/20 | 9/20 | 13/20 | 4/20 |
|  |  | **Cα-RMSD Å** | 1.07 | 1.84 | 2.68 | 1.30 | 1.00 | 1.10 | 11.99 | 1.17 |
|  |  | **L-RMSD Å** | 2.60 | 1.71 | 1.69 | 0.59 | 0.81 | 1.00 | 4.82 | 1.49 |
| Weng et al.<sup>13</sup> | RosettaDock refinement | **Rank<sup>b</sup>** | 3/50/102 | 3/50/25 | 3/50/30 | 8/50/16 | 15/50/24 | 21/50/26 | 2/50/14 | 2/50/15 |
|  |  | **I-RMSD Å** | 0.92 | 4.92 | 2.93 | 1.68 | 1.73 | 1.97 | 0.88 | 1.22 |
|  |  | **L-RMSD Å** | 3.37 | 8.08 | 6.59 | 8.24 | 7.83 | 6.67 | 2.58 | 3.99 |
| Zaidman et al.<sup>14</sup> | DBSCAN-clustering | **Rank<sup>c</sup>** | 2/35 | 1/29 | 2/22 | 71/76 | 20/35 | 18/61 | 3/17 | -<sup>e</sup> |
|  |  | **Cα-RMSD Å** | 2.26 | 0.5 | 2.26 | - | - | - | - | - |
|  |  | **L-RMSD Å** | 1.13 | 3.35 | 1.35 | - | - | - | - | - |
| Liao et al.<sup>12</sup> | HOPAC rescoring | **Rank<sup>d</sup>** | 1/5 | NA<sup>e</sup> | 1/3 | 1/5 | 1/7<sup>f</sup> | NA | 1/5 | NA |
|  |  | **Cα-RMSD Å** | - | - | - | - | - | - | - | - |
|  |  | **L-RMSD Å** | - | - | - | - | - | - | - | - |

<sup>a</sup>Rank of the best MM-GBSA scoring ternary complex over the 20 retained protein-protein docked poses.
<sup>b</sup>Rank of the best solution / the rank of the RosettaDock conformer solutions / the rank of the protein-protein docking poses.
<sup>c</sup>Rank of the best solution over the number of final clusters provided by DBSCAN.
<sup>d</sup>Rank of the best solution is that which corresponds to the best HOPAC score.
<sup>e</sup>Missing values translate to unfound values from the literature source.
<sup>f</sup>Rescore only.

We compared our method with three related studies that reported RMSD and rank outcomes (**Table 1**). The rank, Cα-RMSD, and L-RMSD or I-RMSD (interface root-mean-square deviation) values are presented when available in each study for the complexes constituting our benchmark. A direct comparison with the study of Drummond et al.^5^ was infeasible since the latter methodology produces PROTAC solutions as collective hit-rate – defined as the number of near-native structures within a range of clustered solutions.

Encouragingly, we observed the rank of the crystal-like pose did not exceed 15/20. Additionally, the Cα-RMSD and L-RMSD values were generally below 2Ǻ with the exception of 5T35, 6BOY and 6SIS. Overall, our workflow showed competitive performances in terms of Cα-RMSD with published methods. Just in case of the macrocyclic PROTAC-1 structure (PDB 6SIS) the performance was significantly lower. Nevertheless, with respect to the PROTAC molecule (L-RMSD), our pipeline achieved superior results in most instances.

The compared methods resemble our pipeline in modeling the ternary complex starting from PROTAC building blocks especially when it comes to resorting to protein-protein docking in order to obtain ternary complex poses. However, these methods require constrained templates of known PROTAC confirmation^13,14^ or molecular dynamics simulation^12^. In contrast, our “sum-of-parts” integrative approach^39^ does not rely on any *a priori* knowledge of ternary complexes, although it may benefit from refinement particularly at the protein-protein stage, as observed from the crystal recapitulation results (**Figure 2** and **Figure S1**), and at the linker selection step.

### Validation of the modelling workflow

Considering that the benchmarking of the ICM^27^, which was used to dock POI and E3 cognate ligands, was evaluated in previous studies^40^, we focused on other stages along the pipeline. Specifically, at the first linker stage, following ligation and minimization of the POI^pLig-Linker^ complex, the RMSD was computed between the POI^pLig^ and POI^pLig-Linker^. In doing so, we sought to probe the POI^pLig^ conformer changes upon attachment to the linker. Linkers are reported to play a compensatory role in creating new interactions with the protein that might often stabilize the cognate ligand^10,41,42^ and participate in enhancing the cooperativity between the POI and the E3 proteins^10^. Our experiments show that the linker confirmation with the lowest RMSD changes incurred on the POI ligand conformer are desirable and often result in crystal-like recapitulation of the ternary complex (**Figure S2**).

The performance of the protein-protein docking on ternary complexes using ProPOSE software^43^ was evaluated primarily on the known crystal structures as detailed previously, suggesting that native crystal poses usually resided within the top 15 poses resulting from protein-protein docking (**Table 1** and **Figure S3**). Thus, 20 ternary complexes were retained for each complex from the protein-protein docking step.

Finally, since MM-GBSA scoring was used to minimize and rank the modelled ternary complexes, we mapped the delta Gibbs Free Energy (dG) values corresponding to labelled active and inactive PROTACs. By taking the average dG across the 20 ternary complexes conformations, we observed no separation between the inactive and actives classes – −146.66 kcal/mol (35.91 s.t.d.) for active cases and −145.12 kcal/mol (35.19 s.t.d.) for the inactive cases. As a result, for the purpose of training the SE(3) network on a single conformation from each entry, we resorted to selecting the ternary complex corresponding to the best protein-protein conformation that was predicted using the open-source tool GNN-DOVE^44^ (see **Methods**)

### Prediction of PROTAC activities from modeled structures

Leveraging the modeled ternary complexes, we developed a GCN classifier implemented with an SE(3)-tansformer^32^, whose roto-translational equivariance substantially improves the model’s efficiency. This feature enables the model to generalize effectively from a smaller training dataset by leveraging learned knowledge across diverse spatial configurations of ternary complexes^32,45^. Further, a geometric deep learning model is more suitable to glean nuances surrounding PROTACs design including the presence of y-lysine, sumoylation, and terminal N-terminus amine which are crucial for the UBS degradation through E3-ligases^46^. Our SE(3)-transformer classifier was trained using 3-D descriptors from atomic node embeddings, covalent bond matrices, and 3-D coordinates that were used to compute pairwise distances (more details in the **Methods**). In addition, we concatenated 2,048-bit Morgan fingerprints (MFPs) and 200 RDKit-generated features^47^ to the graph learned representation, in order to enrich the representation of the PROTAC macromolecule^48^.

The general network architecture of the SE(3)-transformer classifier is illustrated in **Figure 3**. Initially, the nodes and edges are embedded and are used to build a graph representation. The graph is then fed into the SE(3)-transformer to predict three features per atom. The resulting atomic features are then convoluted over in a temporal convolutional neural network (TCN) to reduce dimension down to one feature per atom. In turn, adaptive average pooling followed by attention layers and a series of fully connected layers (FCNs) are applied to obtain the classification prediction.

**Figure 3.**
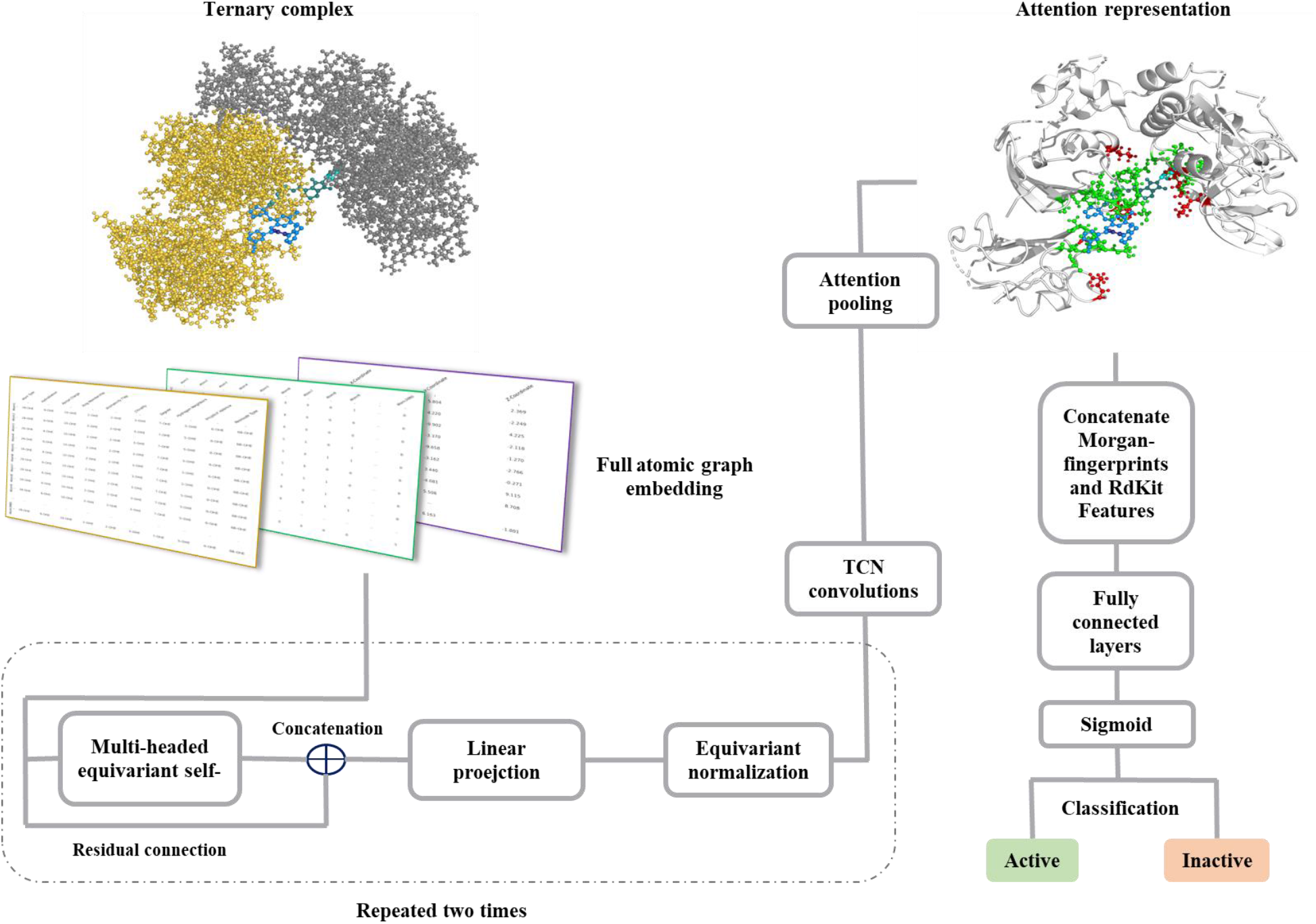
SE(3)-transformer component of PROTACable. The scoring function used is a trained SE(3)-transformer network which predicts on full atomic graph given structures of the ternary complex.

Overall, our model achieved an AUC-ROC of 0.92, an accuracy of 0.86, and an F1-score of 0.89, demonstrating high accuracy in predicting PROTAC activity for computationally modelled ternary complexes (**Figure 4a**).

**Figure 4.**
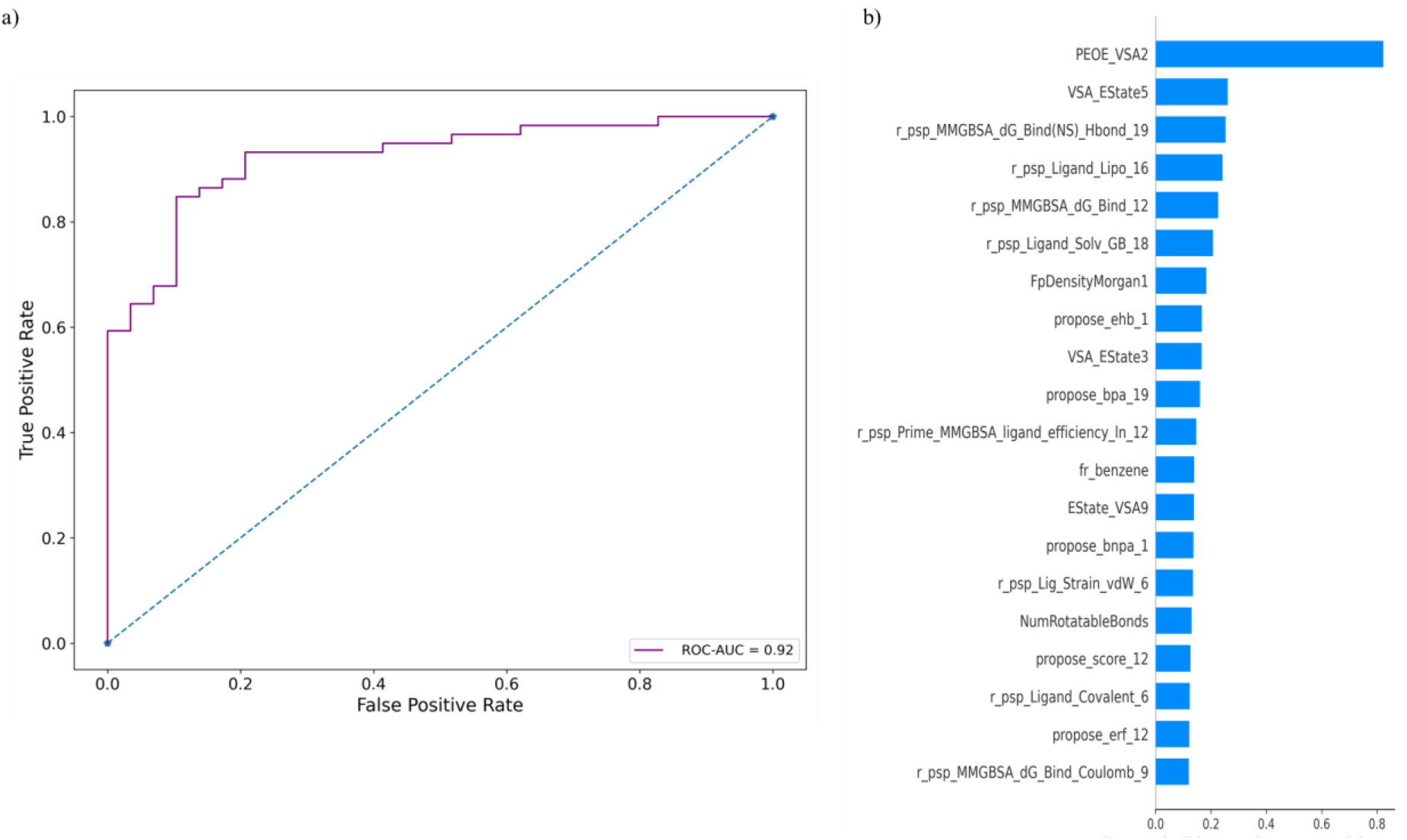
a) The probabilities for each class (positive/1 or negative/0) were used to plot the ROC-AUC curve where the green line represents the SE(3) equivariant GCN performance. SE(3)-transformer classifier achieved an AUC-ROC of 0.92. **b)** The SHAP plot reflects the relative feature contribution (scaled) of each variable to the prediction outcome from the CatBoost model – higher horizontal bars reflect greater feature weight.

### Feature Importance in PROTAC design

In addition to proposing a trained SE(3) network to classify PROTAC activity, we leveraged the merged dataset from PROTACpedia^31^ and our own data resulting from the modelled ternary complexes to shed light on features predictive of PROTACs activity. Since PROTACs are commonly characterized for exceeding the ‘rule of five’^42^ in addition to their dependence on other factors such as cellular availability of the addressed E3 ligase^10^, feature importance may further guide selection of PROTAC from *in silico* modelling and help elucidating structure-activity relationships which govern PROTAC activity^49^.

The merged dataset used in this analysis included scores from small-molecule docking, RMSD changes upon linker ligation, protein-protein docking and interaction scores, MM-GBSA rescoring, and various energy properties. In order to complement the feature descriptors, we also computed 2,048-bit MFPs and 200 RDKit features of the PROTAC, covering properties like hydrogen bond donors and acceptors, TPSA, MW, and fraction of sp3 atoms^47^. We also factored in the averaged and summed features from the 20 protein-protein docking conformations for each entry.

Thus, we deployed a CatBoost classifier that is distinct from the main SE(3)-transformer model. The fine-tuned CatBoost classified helped in the elucidation of the importance of each feature, as detailed in Methods and illustrated in **Figure 4b** via SHAP method^50^.

Interestingly, we observed that the feature with most significance were associated with PEOE_VSA2 and VSA_EState. These 2-D topological/topochemical attributes encode the atomic contributions influenced by partial total charge (PEOE) and molar refractivity to the Van der Waals surface area (VSA)^51^. Components such as MMGBSA_dG_Bind_HBond, MMGBSA_dG_Bind_vdW, and r_psps_Ligand_Lipo, connected to MMGBSA scores, emerged as notable among the SHAP values, reflecting the free energy of binding with emphasis on hydrogen bond energies, vdW interactions, and lipophilicity properties.

For protein-protein docking (ProPOSE), comparable feature weights were attributed to protein-protein electrostatic reaction field energy (Eerf) and protein-protein polar buried surface area (Ebpa)^43^. Descriptors of hydrophobicity appear to be important as well and are in line with PROTAC structures that are abundant in hydrophobic functional groups (e.g., fr_benzene).

Notably, hydrophobicity may contribute to binding conformations that occlude solvent from the interacting proteins, underscoring the critical trinity of POI-E3 ligase protein compatibility and induced-fit binding of PROTACs. This is further supported by metrics like ligand efficiency (r_psp_Prime_MMGBSA_ligand_efficiency)^52^.

In addition, the emphasis on Hbond features seen in the plot suggests an important role of both inter and intramolecular hydrogen bonding or IMHBs, considering PROTAC is a macromolecule^49,53,54^. These descriptors, coupled with lipophilicity tendencies, might have played pivotal roles in determining PROTAC activity by affecting cell permeability and target binding proficiency of given PROTACs^49,53,54^. Conversely, the alterations in Hbond post-minimization may represent values suggestive of enhanced binding affinity to the cognate protein^55^. We hypothesize that a feature like NumRotatableBonds implies a compact PROTAC molecule, maintaining stable binding affinity towards either cognate POI or E3 ligase.

Overall, by decomposing the energy contributions from each stage, we identified critical features such as hydrophobicity, Hbond characteristics, and lipophilicity tendencies as key determinants in PROTACs’ design and optimization.

### PROTACable: a pipeline for de novo design of PROTACs

We developed an automated platform for the de-novo design of PROTACs, tailored for new POI and cognate ligands. The herein suggested platform is based on the primary workflow discussed above. However, to enhance accessibility, most commercial dependencies were replaced with open-source alternatives.

The general process of the platform is outlined in **Figure 5** and is divided into five stages in the GitHub repository:

**Figure 5.**
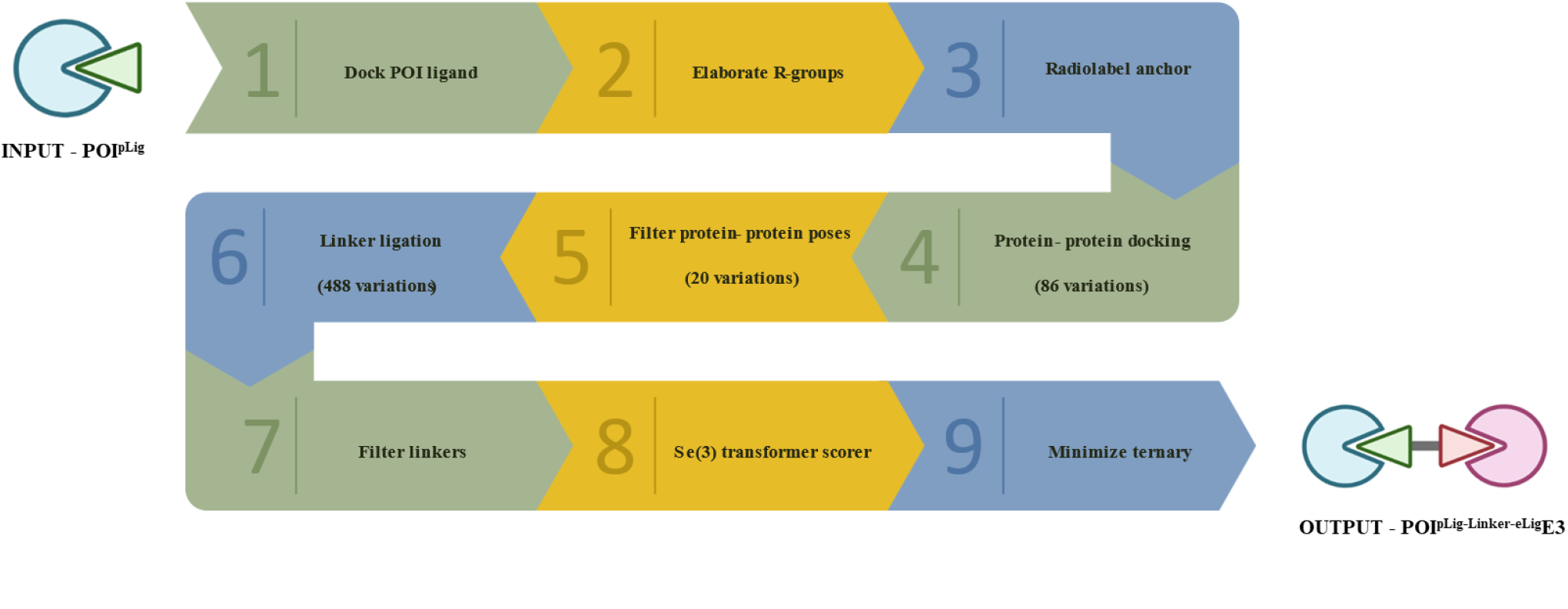
The PROTACable pipeline: the input is a POI with or without a pre-defined ligand pose. A series of steps depicted are the internal working of the five stages of the proposed modelling process and involve the usage of the curated E3 and linker library. The output is a range of up to 20 modelled candidate ternary complexes.

– Stage I – Docking POI and Elaborating Variations:

Given a starting POI PDB file and an SDF ligand file, the user may opt for using the incorporated GNINA^56^ engine for docking or use a predefined confirmation. Next, a preset of amide and carboxyl R-group structures are attached to the POI^Lig^ user-specified anchor – the linker attachment point. This yields more realistic and synthetically accessible PROTACs^41^. Finally, the POI^Lig^ must be updated for the new anchor atom.

– Stage II – POI-E3 Docking and Pose Filtering:

A pre-modelled diverse E3 ligase library comprising 86 E3 ligase proteins is docked combinatorically using ProPOSE^43^ and the POI^Lig^ to yield 86 solutions (unique E3^eLig^). Subsequently, the 100 protein-protein pose output are filtered to 20 PROTAC-productive poses.

– Stage III – Linker Ligation and Pose Filtering:

Using the 488 linkers extracted throughout the antecedent modelling pipeline, each of the 86 solutions and their corresponding 20 PROTAC-productive poses are “mixed and matched” combinatorically with each linker. During the process, an extensive conformation search is performed on each linker to exhaust the extent of compatibility for each POI^pLig^-linker pair in taking into account the distance constraint between the POI^pLig^ and E3^eLig^. Consequently, linkers failing to bridge the POI^pLig^ and E3^eLig^ are discarded. The output of this stage is the full ternary complex with occasional abnormally longer bonds between the POI^pLig-Linker^ and E3^Lig^ prompting further optional minimization to adjust the linker ligation step between POI^pLig-Linker^ and E3^eLig^.

– Stage IV – SE(3) Transformer Network Score Prediction:

The results from the previous stage would be pooled in a single output directory. The pre-trained SE(3) transformer model can then be used to predict which of the ternary complexes are likely active or inactive. This stage encompasses the embedding of each complex and the subsequent prediction. The ranking of activity probability for each complex is generated. In addition, the top 20 probabilistically active ternary complexes are saved.

– Stage V – Ternary Complex Minimization:

As aforementioned, an additional minimization step is optional to complete the modelling of the ternary complex. Therefore, we employed Schrödinger’s PrepWizard utility for the multipurpose of specifying the attachment pattern that ensures optimal ligand ligation, ternary structure minimization and fixing any remaining structural anomalies. Alternatively, the user may choose other minimization tools in order to fulfill this requirement.

Upon the consecutive execution of Stage I-V, up to 20 top-scoring ternary complexes will be proposed for potential *in vitro* validation.

### Case study – G9A

The G9a protein, also known as euchromatic histone-lysine N-methyltransferase 2 (EHMT2), plays a crucial role in histone methylation, influencing gene expression and cellular differentiation^57^. Targeting G9a is advantageous in therapeutic strategies, particularly in cancer treatment, due to its significant involvement in tumor growth and metastasis, as evidenced in various studies highlighting its overexpression in cancers like melanoma^57^.

In order to showcase the applicability of PROTACable pipeline, the G9A inhibitor UNC0642 (IC_50_ 1−20 μM in T24, J82, and 5637 cells^58^) (**Figure 6A**) was proposed for developing new PROTACs. The solved crystal structure of a G9A structure with inhibitor UNC0638^59^ was used to validate the docking pose of the closely related analog SMI UNC0642 after Stage I of PROTACable. By applying the full pipeline, more than 1000 PROTAC ternary complexes were produced. Remarkably, all of the top 20 scoring novel PROTACs, which are retained after Stage V, comprised a VHL-based E3 ligase (**Figure 6B**). Despite this, ternary complexes containing different E3 ligases, such as RNF4, emerged when taking the top 50 scored ternary complexes (**Figure S4**).

**Figure 6.**
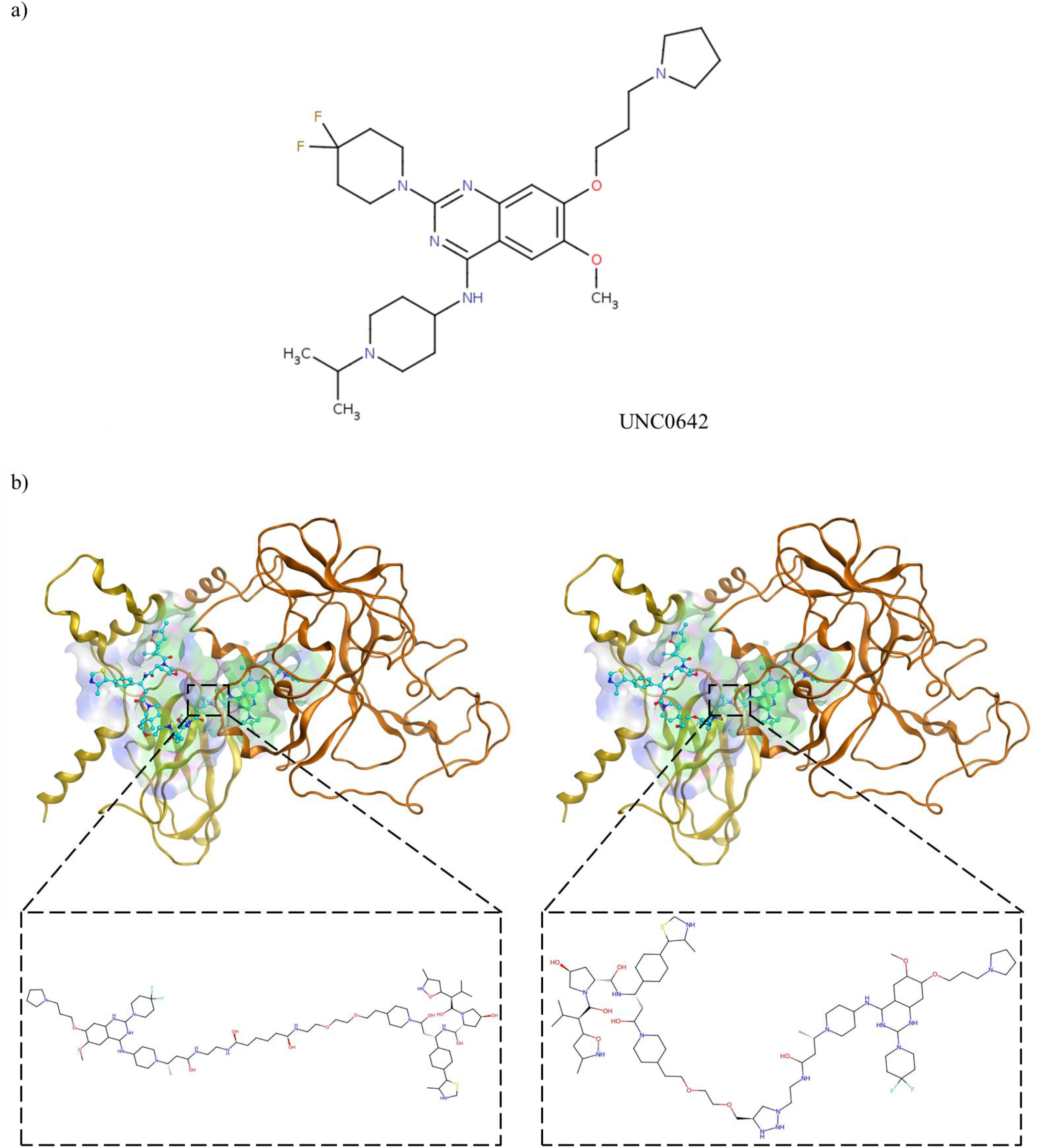
a) G9A inhibitor UNC0642 structure, used as the candidate PROTAC warhead, is shown. **b)** Output of the PROTACable pipeline. The top two scored candidate ternary complexes are shown. In the figure, the pocket surface highlights polar (pink), hydrophobic (green), and solvent-exposed (dark blue) features. The PROTAC is colored in cyan whereas the E3 ligase is colored in gold and the protein of interest (G9A) is colored in dark orange.

Overall, the results obtained from this case study demonstrated the robustness of the PROTACable pipeline in producing novel and diverse PROTACs with predicted *in vitro* potency.

### Conclusion

Herein, we presented a novel strategy for accurate modeling of the PROTAC-induced ternary complexes in a thoroughly automated fashion. The robustness of the initial 3-D modeling approach was primarily evaluated against crystallized ternary complexes. We observed accurate near-native structure recapitulation for the majority of the protein-protein poses as well as the PROTAC ligand poses. Based on the results, we then trained a SE(3)-transformer to classify activities of PROTAC molecules with high accuracy. Additionally, a CatBoost classifier model was used to perform feature importance, where we observed PROTAC-driven compact hydrophobic interactions as the leading feature in describing activity.

To democratize PROTAC design, we incorporated our workflow in PROTACable, a package based entirely on open-source tools, to allow users to model and screen a diverse combinations of different linkers and E3 ligase complexes before they are triaged for activity in cellular assays.

## Material and Methods

### Preparation and cleaning of database

The PROTACpedia database^31^ was used as database. PROTACs were manually deconstructed into ligands and linker (if a linker exists) to populate a non-covalent terminating linker and ligand library, where linker entries were annotated at ligation to guide re-ligation. From this database, 86 unique E3 ligase binders, 262 unique POI binders corresponding to 117 unique Protein Data Bank (PDB) IDs and 664 unique linkers were identified. For POIs with unidentified PDB IDs throughout the database, we manually assigned IDs based on ligand similarity to an existing complex’s ligand within the PDB repository. When multiple IDs for a single PROTAC were found, all of them were kept until they were evaluated at the docking stage. Moreover, PROTACs that exist in the PROTACpedia and that had activity annotation for additional targets were accommodated into their own entries. This resulted in 1,236 entries in total. Each ligand unit from the PROTAC macromolecule was washed, protonated for dominant protomer at pH 7.4 and was converted into 3-D conformer using Molecular Operating Environment (MOE) program^60^.

### Small-molecule molecular docking and linker selection

Internal Coordinate Mechanics software (ICM)^61^ was used to dock each ligand into its respective POI or E3 target. A maximum of one pose was allowed to be generated with an effort of 8.0. When multiple PDBs were available for the same POI, the structure showing the best docking score was retained. Linker docking was similarly done by ICM although the grid was generated while keeping the previously docked ligand as part of it by. Additionally, in the linker case, the effort was set to 1.0 and up to 10 poses were retained to encourage conformational sampling at high speed. Openbabel^62^ was used to convert the output sdf files into PDB and mol2 files. The candidate linker, in the PDB format, was then selected based on the shortest Euclidean distance between the linker PDB and POI ligand anchor points using via a in-house python script utilizing the PDB reader Biopython^63^.

### Ligation and energy minimization

A shell script was developed to reenact bonding between constituent ligands and linker. Briefly, simple rules were devised to trace atoms throughout the pipeline. Automated adjustments are made in the cases where the linkers are nonexistent, are single atoms, or have no second attachment sites on the POI side. Prior and following ligation, a constrained minimization job was performed using the Prepwizard command line utility by Schrödinger using default parameters^64,65^.

### Protein-protein docking

Protein-protein docking was carried out using the software ProPOSE v.1.0.2^43^. AmberTools 21 was used to provide each protein-protein docking run with force field parameters, topology and coordinate files^66^. POI and E3 were prepared as follows:

1. POI and E3 ligands are stripped of ions and salts. The partial charges and formal charges were calculated using Gasteiger-Marsili sigma partial charges implemented in antechamber^43,66^. Finally, the corresponding mol2 force field parameterizing files are generated.
2. Apo forms of POI and E3 are fixed for chain breakage using the python PDBFixer tool^67^ where applicable. Any non-canonical amino acids are substituted for their canonical counterparts (in order to comply with the ff14SB force field dictionary for accepted amino acids^68^). Finally, amber-compatible topology PDB files are generated.
3. Output files from 1) and 2) – prepi and frcmod for the small molecules And amber-compatible PDB for the proteins – are passed to tLEaP tool where the amino acid force field was set to ff14SB, and the small molecules force field was set to gaff. TIP3P was used as a water model.
4. tLEaP-generated inputs were passed into the ambpdb tool – which converts the resulting prmtop and incprd files into a Tripos format (mol2) complex file. Both the final mol2 files and the prmtop files for the POI and the corresponding E3 ligase are used as inputs for the ProPOSE software.

When the input files were obtained, the parameters of the ProPOSE software were setup such that the reference protein was the POI, while the E3 protein was allowed to dock dynamically as a ligand. The number of generated poses was set to 100 poses. 20 poses were filtered down to PROTAC-relevant conformations through the calculation of the average of the sum of the average of the centroids of ligands and the city-square distance between the E3 ligase ligation/anchor point and the POI^pLig-Linker^ calculated as:

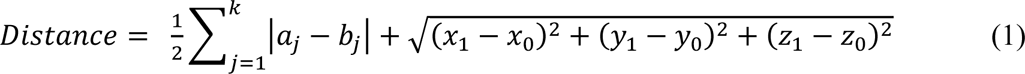

Where *α* is the POI^pLig-Linker^ attachment site for the E3 ligand, *b* is the E3^eLig^ anchor site to the POI^pLig-Linker^; *X*_1_, *Y*_1_, and *Z*_1_ are the POI^pLig-Linker^ average coordinates, and *X*_0_, *Y*_0_, and *Z*_0_ are the E3^eLig^ average coordinates.

### Ternary complex refinement and filtering

POI^pLig-Linker^ and E3^eLig^ output files from ProPose were converted into PDB with OpenBabel^62^, concatenated and unique atom naming was set with pdb4amber^66^. Schrödinger ‘s Prepwizard was used to fix bond orders^69^. The final ligation between the E3^eLig^ and the POI^pLig-Linker^ was performed as previously described.

### Rescoring and optimizing the ternary complex

Using Prime command line tool from Schrödinger^65^, the ternary complex was rescored with the MM-GBSA approach while optimizing the binding site in the presence of the macromolecule.

### Reconstructing crystallized ternary complexes

The procedure described above was used for modeling the crystallized PROTACs found in the PDB repository (PDB IDs: 5T35, 6BN7, 6BOY, 6HAX, 6HAY, 6HR2, 6SIS, 6W7O, 6W8I, 6ZHC, 7JTO, 7JTP, 7KHH, 7PI4, and 7Q2J)^35^. RMSD was calculated using the following tools: Cα-RMSD was calculated using MOE chain pairwise alignment tool^60^, whereas the PROTAC L-RMSD was calculated using OpenBabel command line tool^62^.

### SE(3) transformer network

The dataset (1,236 rows) was scaled up to oversample minority inactive class to yield a ratio of 825: 616 active to inactive class and was split into 80-12-8 train-val-test sets taking into account butina clustering with Tanimoto threshold of similarity set at 60%. The resulting held-out test set comprised a total of 88 samples. The features include 2,048 bit morgan fingerprints and the 200 ‘rdkit2D’ normalized features using Descriptastorus library^47^.

Despite obtaining 20 poses for each row, one candidate pose was retained using GNN-DOVE^44^. In GNN-DOVE^44^, the mode was set to 1 for many-protein scoring paradigm and −1-fold was used. The POI-PROTAC-E3 interfaces (comprising a maximum of 400 protein residues) were extracted using BioPython^63^. Subsequently the local and global structures of the ternary pocket are accounted for by a neighborhood aggregation scheme and a K-Nearest Neighbors (KNN) graph representation, similar to previous implementations^70^.

Nodes were characterized by features such as atom type, associated amino acid type, hybridization, partial charge, ring membership, aromaticity, chirality, formal charge, number of neighboring heavy atoms, total number of hydrogens (implicit and explicit), implicit valence, and 3-D coordinates (120 features per atom). Moreover, the node’s position in a residue-specific local coordinate system was calculated, based on the principal axes of the residue’s atomic structure and using techniques such as the Rodrigues’ rotation formula (Equation 2) to ensure proper alignment with the global coordinate system.

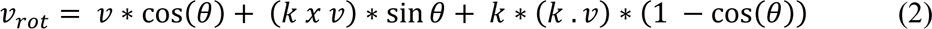

where v is the vector to be rotated, k is the rotation axis, theta is the angle of rotation.

Edges include a distance-based feature and a covalent bond feature for each i-j (where I and j are covalently bonded atoms). Furthermore, a set of relative position for protein atoms (atom – Cα atom) and ligand (atom – ligand fragment atom) are calculated. These features provide a nuanced view of the spatial relationships between atoms, leveraging the calculated local coordinate systems for each residue^45,71–73^. Both the node and edge features were linearly embedded into a 32-dimension tensor before constructing the graph. The graph was constructed based on KNN of 30 nearest atom around each atom with x1.5 weight given to ligand-ligand edges and x3.0 given to ligand-protein edges while embedding.

The model is based on two SE3 equivariant attention blocks, followed by a TCN block, a multiheaded attention block^58^, an attention pooling block, three fully connected layers, and a classifier. The model is implemented using PyTorch Lightning and the Deep Graph Library^74,75^. The model was initialized with Kaiming He uniform distribution weights^76^. Binary cross entropy with logits (BCEWithLogits) was used as the loss function while Adam was used as the optimizer set with a learning rate of 1E-04. The model was trained for 50 epochs.

### Feature importance

CatBoost Classifier^77^ was used for computing weights used in feature importance. Identical training, validation and testing sets used in the SE(3) transformer were used with the CatBoost classifier for elucidating feature importance. The classifier were fine-tuned for optimal hyperparameters based on stochastic random grid search and 3-fold cross validation.

For plotting feature importance, an off-the shelf implementation of SHapley Additive exPlanations (SHAP)^50^ algorithm was used.

### Metrics

The accuracy of predictions was calculated using the following formula:

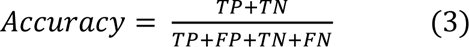

The ROC curve was plotted using true positive rate (TPR) and false positive rate (FPR):

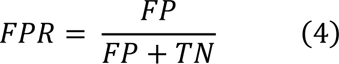

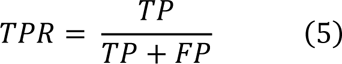

The false positives and the false negatives were also evaluated using F1 scores – computed according to the formulas presented below:

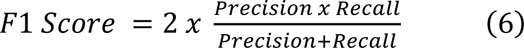

### Linker ligation in PROTACable pipeline

Up to 500 conformers are generated for each of the 488 linkers to exhaust torsional energy profiles and rotatable for birding the POI^pLig^ and linker. A score of intramolecular and intermolecular steric clash and distance between the POI^pLig^ and E3^eLig^ is minimized iteratively for each possible conformer. The linker candidate with the best score is further filtered out if the distance between POI^pLig-Linker^ and E3^eLig^ is greater than 5 Ångstroms. Radiolabels for proximity calculations were employed as previously detailed. All implementations were carried out with the RdKit^78^ library.

### Open-source software in the PROTACable pipeline

For small molecule docking, GNINA v1.0^56^ was used. The configuration of the GNINA docking protocol was preset to an exhaustiveness of 16.

### Computational resources

The modeling of the three-dimensional PROTACpedia dataset was run on 20 x Intel(R) Xeon(R) Gold 6130 CPU @ 2.10GHz. The SE(3)-transformer was trained using 4 Tesla V100 GPUs. All calculations were performed on the Juggernaut computing cluster at the Vancouver Prostate Centre.

## Data availability

The 3-D modelled dataset and PROTACable pipeline is available at: https://github.com/giaguaro/PROTACable/

## Author Information

### Corresponding Author

**Artem Cherkasov** – Department of Urologic Sciences, Faculty of Medicine at the University of British Columbia, 2660 Oak St, Vancouver, BC V6H 3Z6. ORCID: https://orcid.org/0000-0002-1599-1439.

## Acknowledgement

We extend our heartfelt thanks to Dr. Mani Larijani, Justin King, and David Huebert for their invaluable intellectual contributions to this manuscript. Their insightful conversations and expert perspectives have significantly enriched our work, especially in prioritizing oncological targets. We are deeply grateful for their expertise, thoughtful feedback, and collaborative spirit, which have been instrumental in enhancing the quality and depth of our PROTAC research.

## Notes

The authors declare no competing financial interest.

