## Supplemental figure 1 for "PROTACable is an Integrative Computational Pipeline of 3-D Modeling and Deep Learning to Automate the De Novo Design of PROTACs"

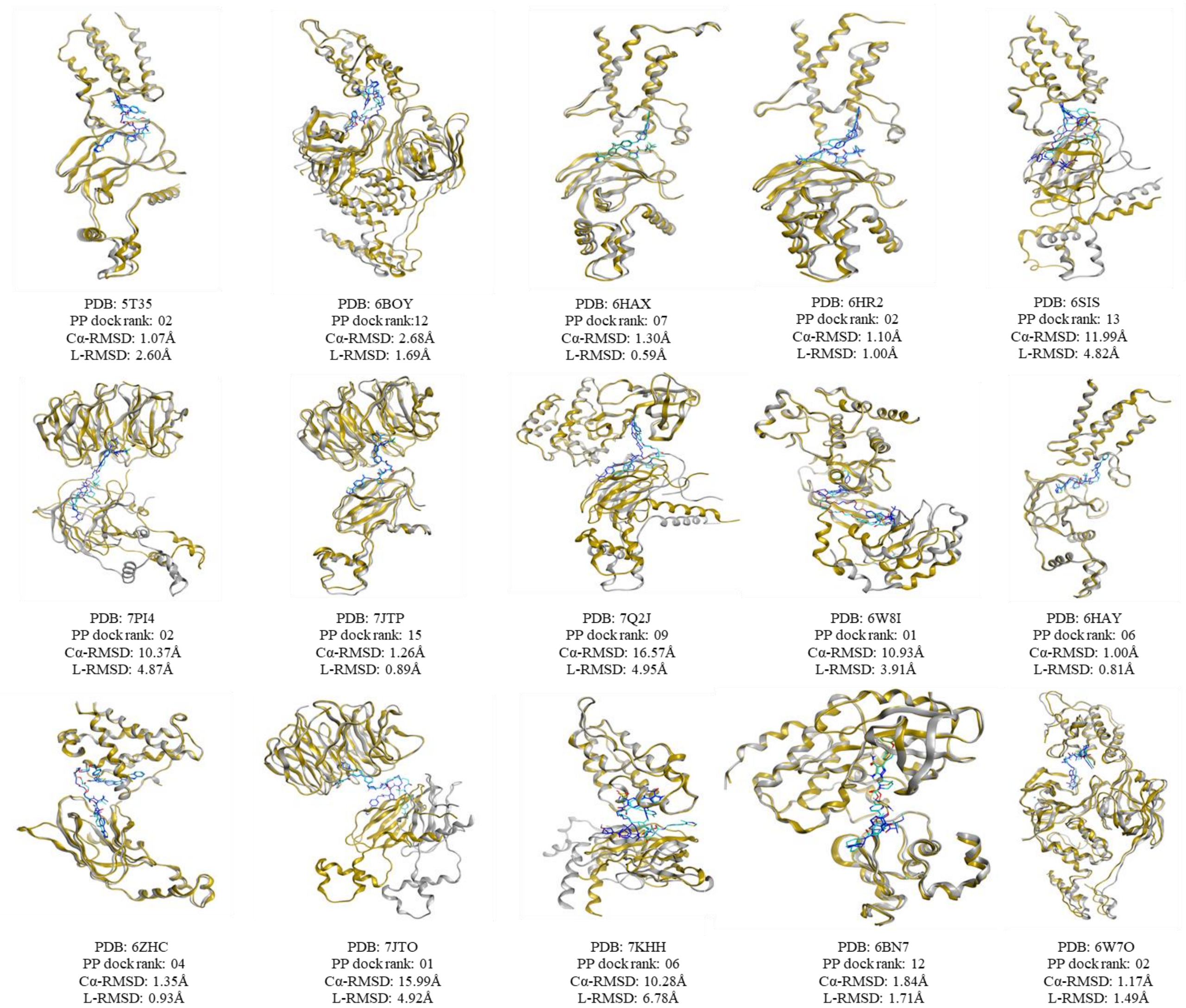


**Figure S1.** Benchmarking of the pipeline on crystallized PDB entries. Crystal ternary proteins are colored in grey while respective PROTAC poses are colored in blue. Modelled ternary proteins are colored in gold while respective modelled PROTAC poses are colored in cyan. Below each complex, the solution rank (out of 20) is shown as well as the Cα-RMSD and the PROTAC L-RMSD between the modeled and the experimental structure.
