## Supplemental figure 2 for "PROTACable is an Integrative Computational Pipeline of 3-D Modeling and Deep Learning to Automate the De Novo Design of PROTACs"

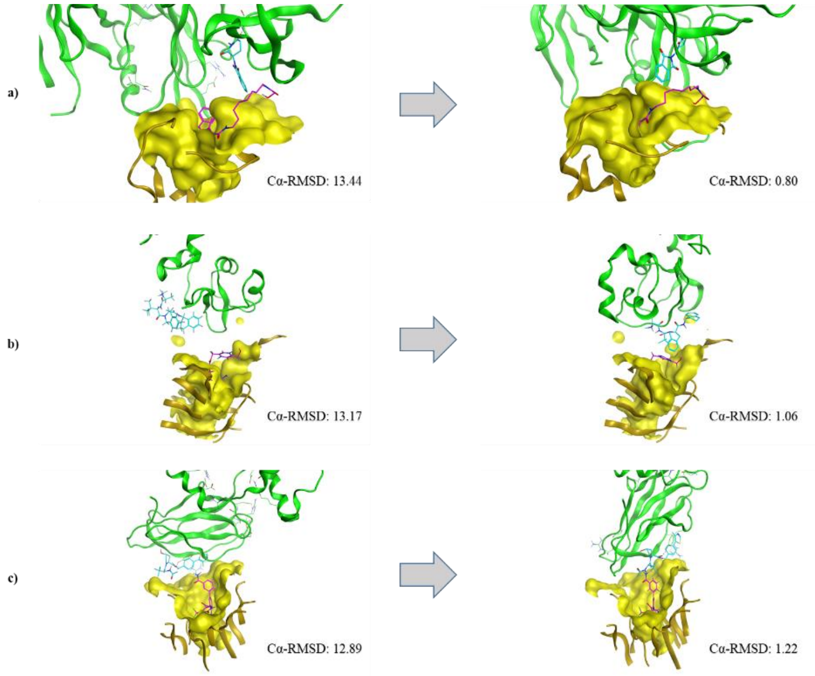


**Figure S2.** Exploring alternate linker confirmations for **a)** 6BN7 **b)** 6W70 and **c)** 7JTP. Shown on the left-hand side are the original conformations resulting from random linker conformation with the protein-protein docking outcome protein's backbone alpha-carbon RMSD (Cα-RMSD) and on the right-hand side are linker conformations with minimal $POI^{Lig}$ RMSD alongside the resulting protein-protein docking output and the respective protein's Cα-RMSD.
