## Supplemental figure 3 for "PROTACable is an Integrative Computational Pipeline of 3-D Modeling and Deep Learning to Automate the De Novo Design of PROTACs"

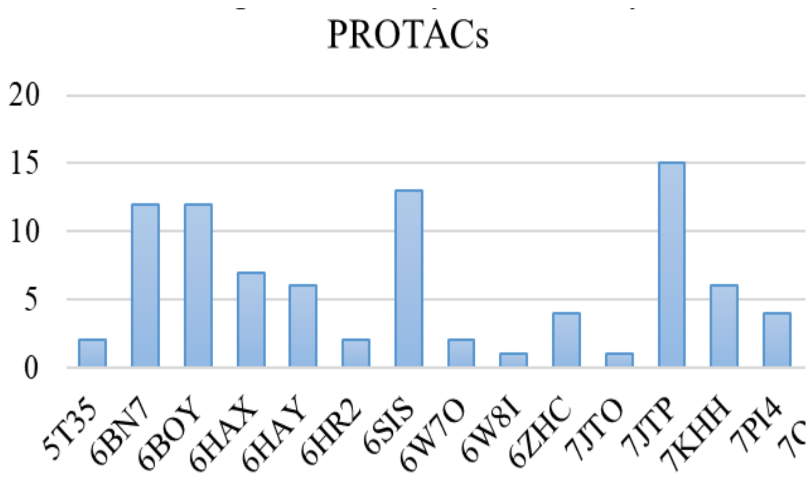


**Figure S3.** The rank of all benchmarked crystals identified through the proposed filtering algorithm, which returns 20 solutions.
