## Supplemental figure 4 for "PROTACable is an Integrative Computational Pipeline of 3-D Modeling and Deep Learning to Automate the De Novo Design of PROTACs"

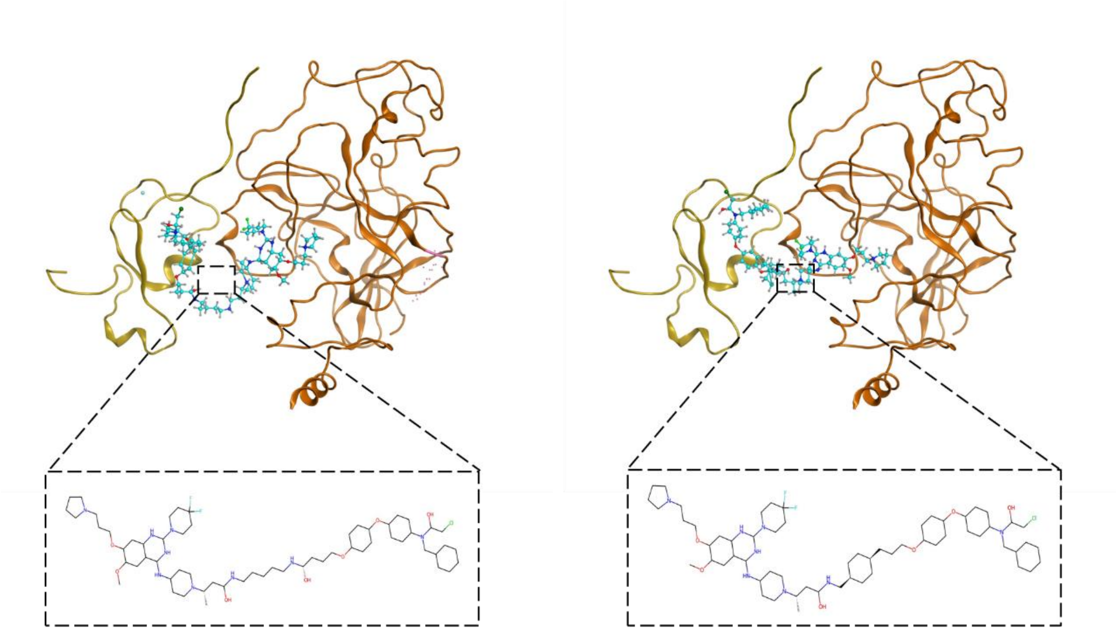

**Figure S4.** Output of the PROTACable pipeline. Two representative examples showcasing diversity of E3 ligase solutions included RNF4 E3 in the top 50 scored ternary complexes. The PROTAC is colored in cyan whereas the E3 ligase is colored in gold and the protein of interest (G9A) is colored in dark orange.
